# Mini-bacterioferritins: Structural insight into a new type of ferritin-like protein from an anaerobic methane-oxidising archaeon

**DOI:** 10.1101/2025.06.06.658376

**Authors:** Martijn Wissink, Sylvain Engilberge, Pedro Leão, Robert S. Jansen, Mike S. M. Jetten, Antoine Royant, Cornelia U. Welte, Tristan Wagner

## Abstract

Ferritins are ubiquitous among life forms, as they are essential for iron homeostasis. Here, we unveiled a novel member of the ferritin family, baptised mini-bacterioferritin. The characterised mini-bacterioferritin was isolated from a microbial enrichment dominated by the methanotrophic archaeon ‘*Candidatus* Methanoperedens’ BLZ2. Its atomic resolution crystal structure reveals a 12-mer assembly with a diiron ferroxidase centre located within a four-helix bundle. Redox cycling experiments on protein crystals reveal a shift in iron position at the active site, which follows the established ferritin catalytic cycle. The 12-mer sphere-like structure harboured six Fe-coproporphyrin III ligands, positioned at the interdimeric interface, a characteristic previously only found in 24-mer bacterioferritins. Phylogenetics, together with structure predictions of closely related proteins, revealed that mini-bacterioferritins form a new clade within the ferritin family that might conserve ancestral traits. Future research will need to investigate the physiological roles of these enzymes, which were unsuspectingly widely distributed among prokaryotes.

**Teaser:** A novel ferritin family, the heme-binding mini-bacterioferritin, assembles as a dodecamer and is widely distributed among microbes.

## Introduction

Iron is essential for life, yet iron homeostasis is challenging due to the low solubility of Fe³⁺ and the toxicity of Fe²⁺ (1). To address these challenges, all domains of life rely on proteins belonging to the ferritin family that store iron in a non-harmful way inside the cell (2). The ferritin family consists of four groups: ferritins, bacterioferritins, ‘DNA-binding proteins from starved cells’ (Dps), and Dps-like proteins (DpsL) (3,4). The defining feature of ferritin-like proteins is the conserved four-helix bundle domain, with helices A and B antiparallel to C and D, connected by a long loop between helices B and C (5). Despite high structural similarities within ferritin-like proteins, amino acid sequences vary, with similarities as low as 20% (6,7).

Ferritins oligomerise as 24-mer sphere-like structures (2,3,7–9). Each subunit contains a nuclear diiron ferroxidase centre that oxidises Fe²⁺ to Fe³⁺ using either oxygen or hydrogen peroxide (10). The iron is then mineralised and stored inside the hollow spherical cavity of the protein, which can hold up to 4,500 iron atoms (11). When iron is scarce, the stored iron is released from the ferritin to the cytosol for incorporation into iron-containing proteins.

24-mer bacterioferritins, found exclusively in prokaryotes, are structurally and functionally similar to ferritins (5,6). The primary distinction is that bacterioferritins bind 12 hemes, which are believed to act as electron conduits, facilitating the reduction of Fe³⁺ within the protein via bacterioferritin-associated ferredoxin (Bfd) (12). Typically, bacterioferritins contain heme *b*, although there is one example of a bacterioferritin from *Desulfovibrio desulfuricans (Dd-*Bfr) that contains a Fe-coproporphyrin-III (coproheme) instead (13,14).

’DNA-binding proteins from starved cells’ (Dps) differ from ferritins and bacterioferritins in several ways, but most notably by forming 12-mer complexes instead of 24-mer complexes (15). As in the case of (bacterio)ferritins, Dps can store iron, and their primary role is to defend cells against a wide range of stresses, including oxidative stress, metal toxicity, thermal stress, and starvation (16). Apart from ferroxidase activity, Dps exhibit additional properties such as DNA binding, endonuclease activity, and protease activity (17,18). Unlike ferritins and bacterioferritins, Dps have their active site at the interface between two subunits, where two iron atoms are coordinated by ligands from both subunits (19).

The most recently characterised group within the ferritin family is the Dps-like (DpsL) group. Structurally similar to Dps, DpsL also form 12-mer assemblies, but its ferroxidase active site more closely resembles that of ferritins and bacterioferritins (20–22). Phylogenetically, DpsL are more closely related to bacterioferritins than to Dps (3,4). Like Dps, DpsL are regulated upon oxidative stress and often associated with additional functions, such as endonuclease activity (4,21).

To hunt for unexplored new ferritin systems used in microbial life, we looked for ferritin-like proteins with sequence identity divergent from known homologues and focused on ‘*Candidatus* Methanoperedens sp.’ BLZ2 (23). The genome of this archaeon harbours five ferritin-like proteins (Supplementary Table S1, Supplementary Figure S1), and two of them present naturally high expression as shown by a complexome profiling study (24). ‘*Ca*. Methanoperedens sp.’ BLZ2 is an anaerobic methane-oxidising archaeon that thrives in anoxic environments, in particular freshwater sediments, where it contributes to the biogeochemical methane cycle by coupling the oxidation of methane to the reduction of nitrate and extracellular electron acceptors such as iron and manganese (25–27). To avoid any biases from recombinant expression (*e.g.*, determining if the protein contains a particular intrinsic cofactor), we sought to natively purify the most abundant ferritin-like protein from this organism.

With this successful approach, we solved the structure of the first representative of a previously unrecognised group within the ferritin family. The atomic resolution structure supported by biochemical studies reveals a 12-mer sphere-like structure harbouring two redox-active catalytic iron per monomer and a total of six coprohemes. Placing the ‘*Ca.* Methanoperedens sp.’ BLZ2 ferritin-like protein in a phylogenetic tree showed the existence of similar proteins that form a distinct clade within the ferritin family.

## Results

### Isolation of mini-bacterioferritin from ‘*Ca.* Methanoperedens sp.’ BLZ2

Anaerobic methane-oxidising archaea (ANME) present significant research challenges due to their lack of axenic cultures, slow growth rates (doubling time ∼30 days in bioreactors), and sensitivity to oxygen. Nevertheless, enrichment cultures of ANME species have enabled the study of their physiology and biochemistry. Here, the enrichment culture of *‘Ca.* Methanoperedens sp.*’* BLZ2 was maintained for over a decade in a bioreactor fed with methane as the sole electron donor and nitrate as the electron acceptor. This setup produced a stable community with 20–40% enrichment of *‘Ca.* Methanoperedens sp.*’* that could be exploited for native purification.

Following a strict anaerobic purification under yellow light, we enriched a ferritin-like protein with an intense pink colour after three chromatography steps. Starting from 17.5 g wet weight biomass, the process yielded approximately 1 mg purified protein. The final purified fraction (with >95% purity confirmed by denaturing polyacrylamide gel electrophoresis, PAGE, Fig. 1A) was identified by mass spectrometry. This protein of 139 amino acids (15.9 kDa) corresponds to a naturally abundant putative bacterioferritin found in meta-proteomics data (24)(Fig. 1). The UV-visible spectrum of the protein confirmed a heme presence, with a distinct Soret peak at 424 nm with a shoulder at 418 nm, and Q bands at 521 and 551 nm resembling the heme spectrum of *Dd*-Bfr (28)(Fig. 1B). Mass spectrometry (MS) of the extracted heme yielded a compound with a m/z of 708.189 [M+H]+, which was predicted to have an elemental composition of C_36_H_36_FeN_4_O_8_ based on MS and MS/MS spectra (Fig. 1C). *In silico* structure prediction using CANOPUS software classified the compound within the porphyrin subclass. Previous work showed that coproheme was detected at m/z 708 using LC-MS (29). Together, these data strongly suggest that the heme extracted from *Mper*-mBfr is a coproheme. Spectral shifts upon air oxidation and dithionite reduction confirmed reversible redox activity, consistent with a coordinated redox-active heme (Fig. 1B).

**Fig. 1.**
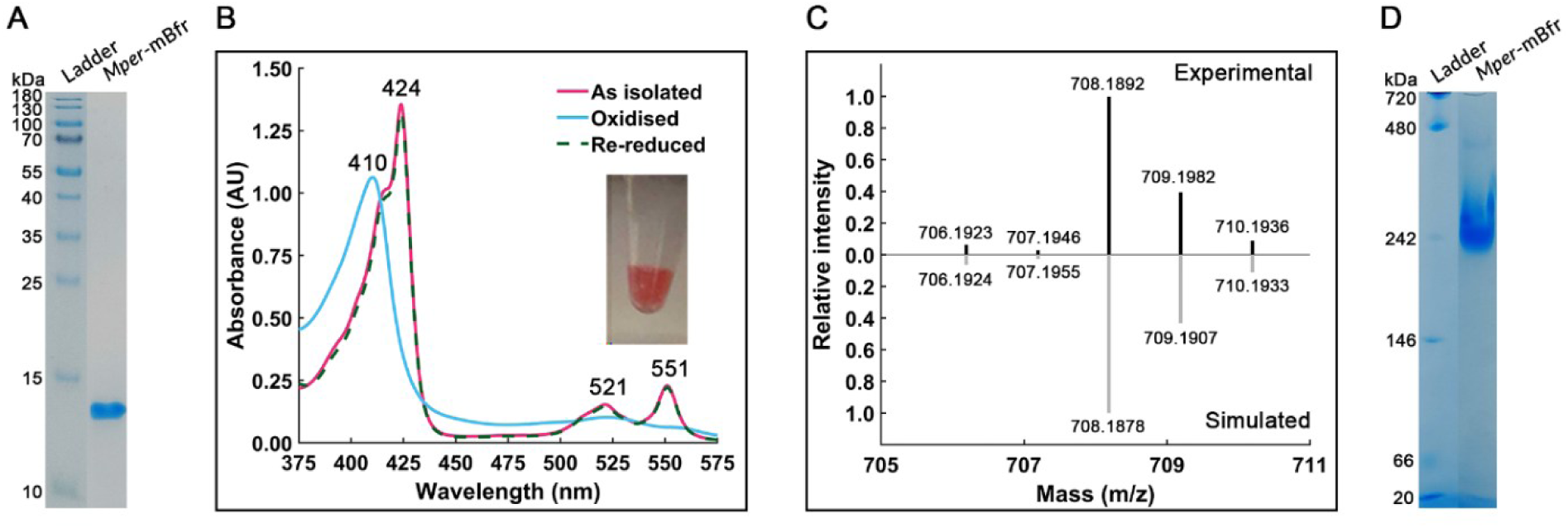
Purification of *Mper*-mBfr. (A) Denaturing PAGE of purified *Mper*-mBfr. (B) UV-Visible spectrum of purified *Mper*-mBfr (2 mg mL^-1^) as isolated, oxidised and re-reduced with 1 mM sodium dithionite with applied baseline correction. The inset shows the protein as isolated under anoxic conditions in an Eppendorf tube. (C) LC-MS spectrum of *Mper*-mBfr extracted hemes versus the simulated MS1 spectrum of C_36_H_36_FeN_4_O_8_. (D) hrCN-PAGE gel of purified *Mper*-mBfr.

The oligomerisation state of the protein was investigated by high-resolution clear native PAGE (hrCN-PAGE) under anoxic conditions. It revealed a protein assembly of approximately 242 kDa (Fig. 1D). While ferritin-family proteins are known to assemble into larger quaternary structures, their size was smaller than expected for a canonical 24-subunit bacterioferritin (*i.e.*, expected to be 381.6 kDa). Based on its origin and characteristics, the protein was provisionally named *Mper*-mBfr (*<u>M</u>ethano<u>per</u>edens* <u>m</u>ini-<u>b</u>acterioferritin). Its unexpected deviation in size and sequence from typical bacterioferritins prompted structural investigation, and anaerobic protein crystallisation was undertaken.

### *Mper*-mBfr forms a 12-mer sphere

*Mper*-mBfr crystallised into bright pink bipyramidal crystals within two weeks. X-ray diffraction data were collected, and the structure was refined to an atomic resolution of 1.07 Å (Table 1). Structural analysis revealed a 12-mer assembly with a 23 tetrahedral symmetry (Fig. 2A), resembling that of Dps (30). In this assembly, the N-terminal ends of three four-helix bundles are facing each other and form the N-terminal pore. Conversely, in the other conformation, the C-terminal ends are facing each other and form the C-terminal pore (Fig. 2A). Since the 432 octahedral symmetry of (bacterio)ferritins only allows for N-terminal-type pores, these pores are also referred to as ferritin-type pores, while C-terminal-type pores are referred to as Dps-type pores (30). Each monomer contains a diiron active site, and one coproheme is bound at the interdimeric interface (Fig. 2).

**Fig. 2.**
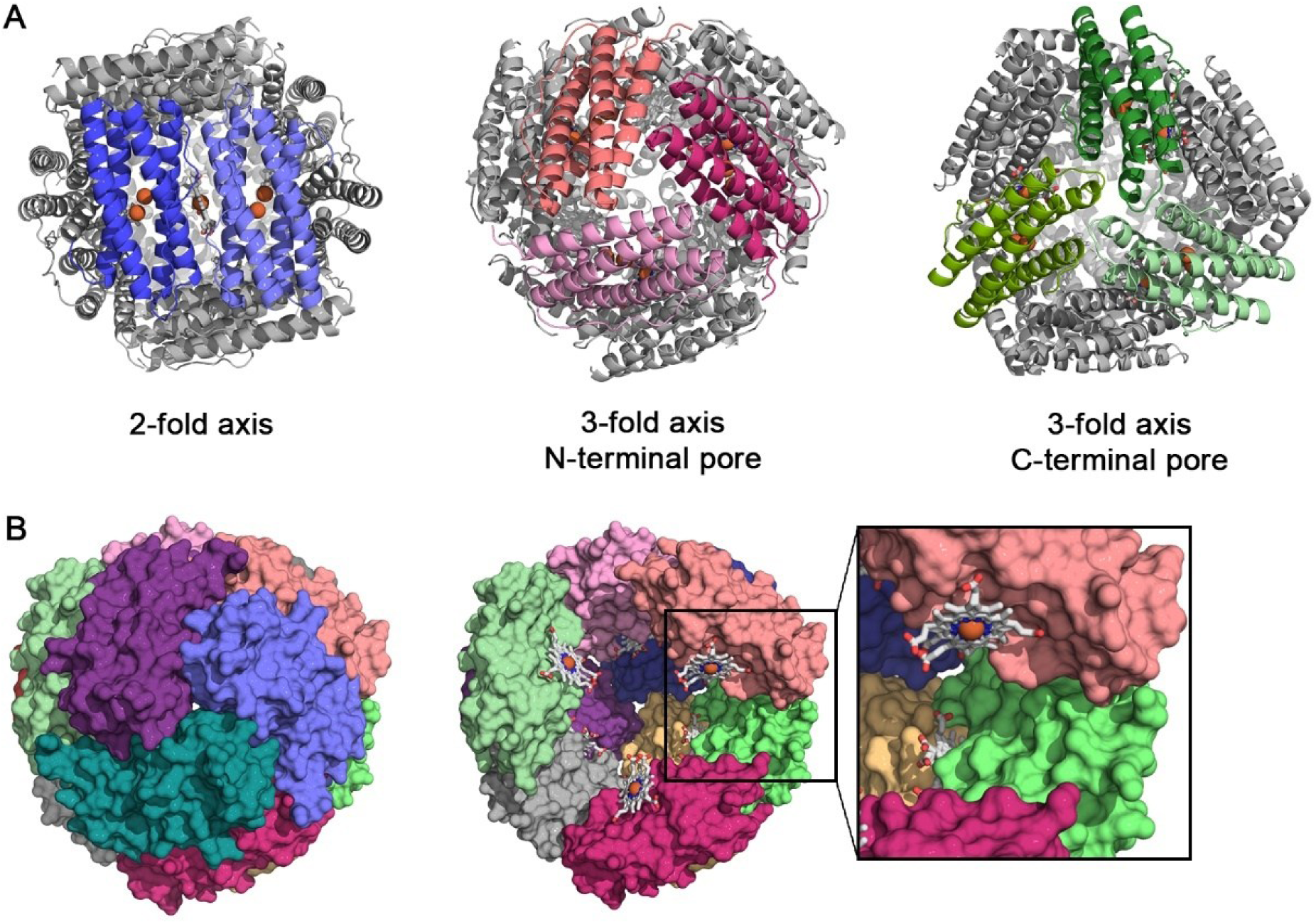
12-mer structure of *Mper*-mBfr. (A) Symmetry interfaces of *Mper*-mBfr are shown as cartoons and viewed along the 2-fold symmetry axis, 3-fold symmetry axis viewing the N-terminal pore, and the 3-fold symmetry axis viewing the C-terminal pore. (B) Surface representation of *Mper*-mBfr (left) and surface view without the front chains (coloured purple, slate blue and deep teal), showing the coordination of heme ligands (right). Heme ligands are represented as balls and sticks, and iron atoms as spheres, with carbons, oxygen, nitrogen and iron coloured as white, red, dark blue, and orange, respectively.

**Table 1:**
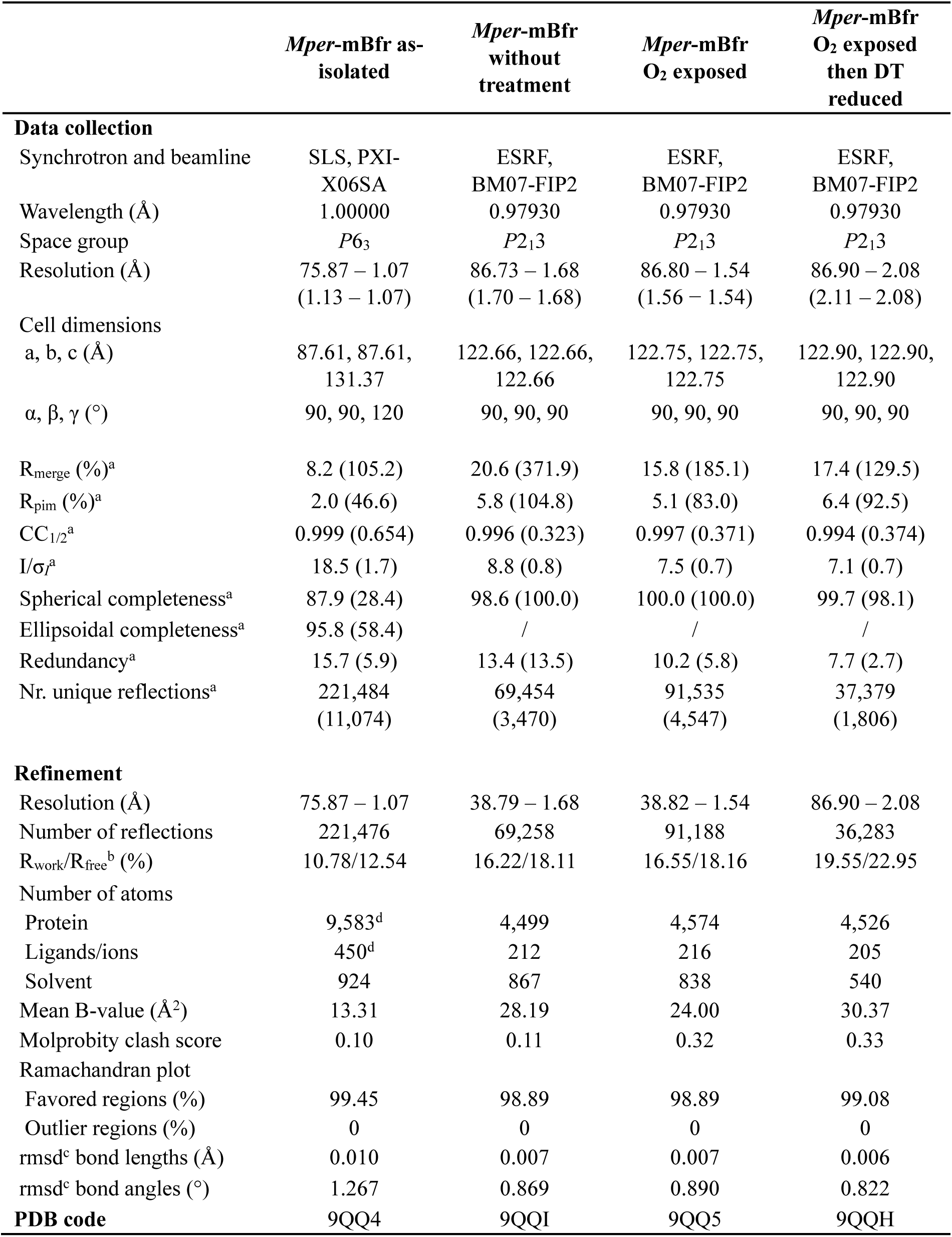
X-ray analysis statistics for *Mper*-mBfr. ^a^Values relative to the highest resolution shell are within parentheses. ^b^Rfree was calculated as the Rwork for 5 % of the reflections that were not included in the refinement. ^c^rmsd, root mean square deviation. ^d^Model contains hydrogens.

*Mper*-mBfr forms a hollow spherical structure with an outer diameter of ∼90 Å and an inner diameter of ∼46 Å (Fig. 2B), with dimensions comparable to *Escherichia coli* (*Ec-*Dps) (∼90 Å and ∼45 Å, respectively) but smaller than *Dd-*Bfr (∼130 Å and ∼85 Å, respectively). The inner volume of *Mper*-mBfr was estimated to be 58 nm³ (see Materials and Methods section for volume calculations), comparable to that of Dps and DpsL, such as *Halobacterium salinarum* Dps (*Hs*-Dps) (56 nm³) (31) and *Pseudomonas aeruginosa* DpsL (*Pa*-DpsL) (75 nm³) (21). In contrast, *Dd*-Bfr has a much larger inner volume of 251 nm³ (13).

### Electrostatic surface potential of *Mper*-mBfr

The outer electrostatic surface potential of ferritin-like proteins plays a crucial role in metal ion transport through pores, and in the case of Dps, in their ability to bind DNA. While the outer charge profile of *Mper*-mBfr reveals interspersed regions of localised positive and negative charges (Fig. 3), the inner surface presents defined positive patches for heme binding, with the rest being negatively charged, creating a suitable environment for storing positively charged iron cations as observed in *Ec*-Dps and *Dd*-Bfr (13,30).

**Fig. 3.**
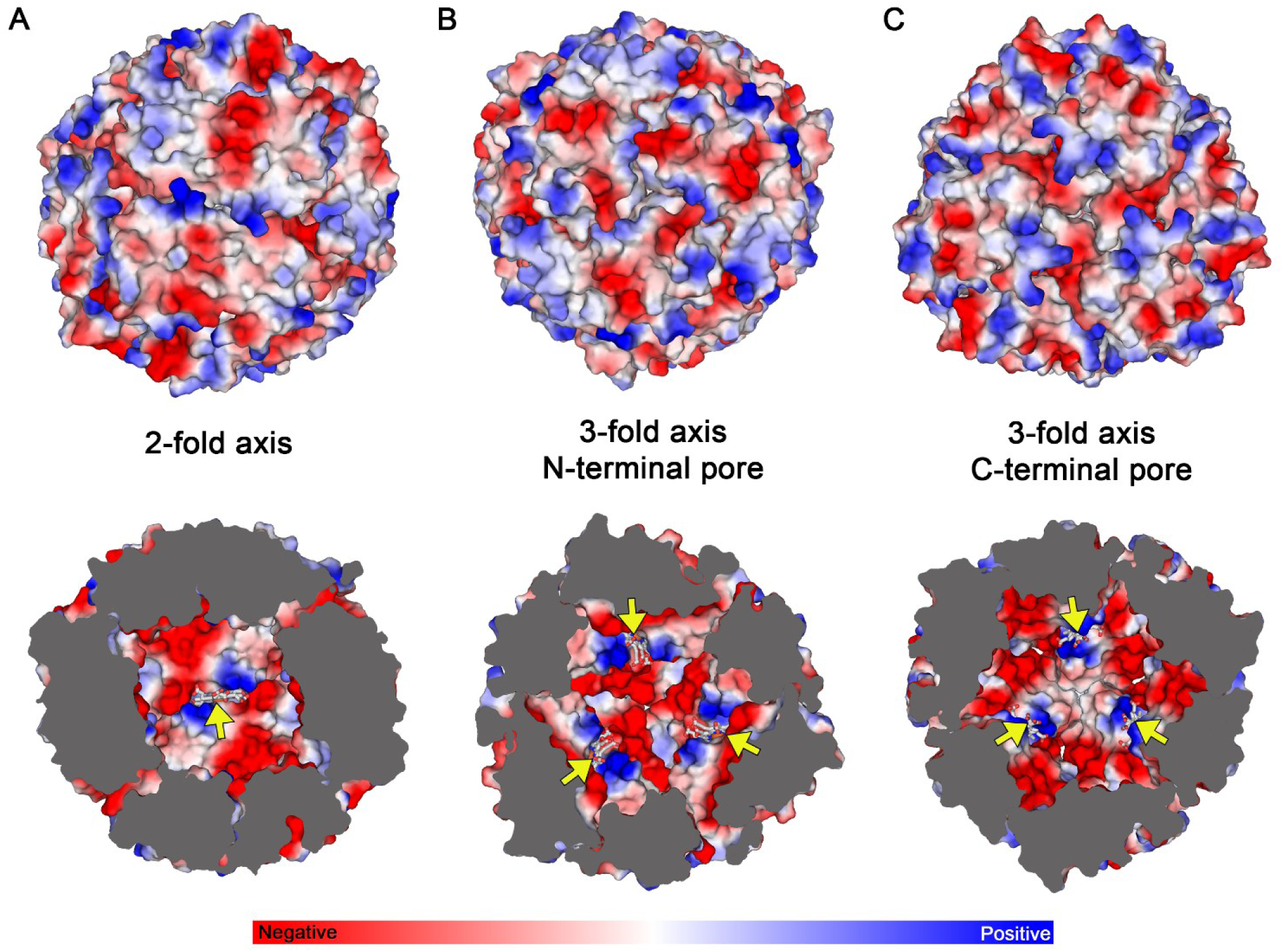
Surface and inner cavity electrostatic charge profile of *Mper*-mBfr. Surface representation of the outer (top) and inner (bottom) electrostatic charge profile along the 2-fold symmetry axis (A), 3-fold symmetry axis from the N-terminal pore view (B) and the 3-fold symmetry axis from the C-terminal pore view (C). Coprohemes are indicated by yellow arrows and displayed as sticks with their iron as spheres, with carbons, oxygen, nitrogen and iron coloured as white, red, dark blue, and orange, respectively.

In ferritin family proteins, three-fold pores have been associated with metal ion transport, typically facilitated by a pathway of negatively charged amino acids (13,20,30,32). In *Ec*-Dps and three types of *Nostoc punctiforme* Dps, the N-terminal pore exhibits a significantly negative potential, while the C-terminal pore remains electrostatically neutral. In contrast, *Mper*-mBfr displays the opposite pattern, with more negative electrostatic potential around the C-terminal pore and a relatively neutral N-terminal pore. This electrostatic distribution resembles what has been observed in *Saccharolobus solfataricus* DpsL and *Dd*-Bfr, suggesting that the C-terminal pore is the more likely route for iron transport into the protein cavity.

### Dual conformation of coprohemes at the dimeric interface

The coproheme is axially coordinated by Met50 located at the dimeric interface along the 2-fold symmetry axis. The atomic resolution electron density map revealed a dual conformation of the coproheme ligand, with each orientation present at approximately 50% occupancy (Fig. 4A). The same dual coproheme-binding alternative conformation has been reported in the coproheme binding *Dd*-Bfr, but also for *Rhodobacter capsulatus* bacterioferritin that binds heme *b* (13,33).

**Fig. 4.**
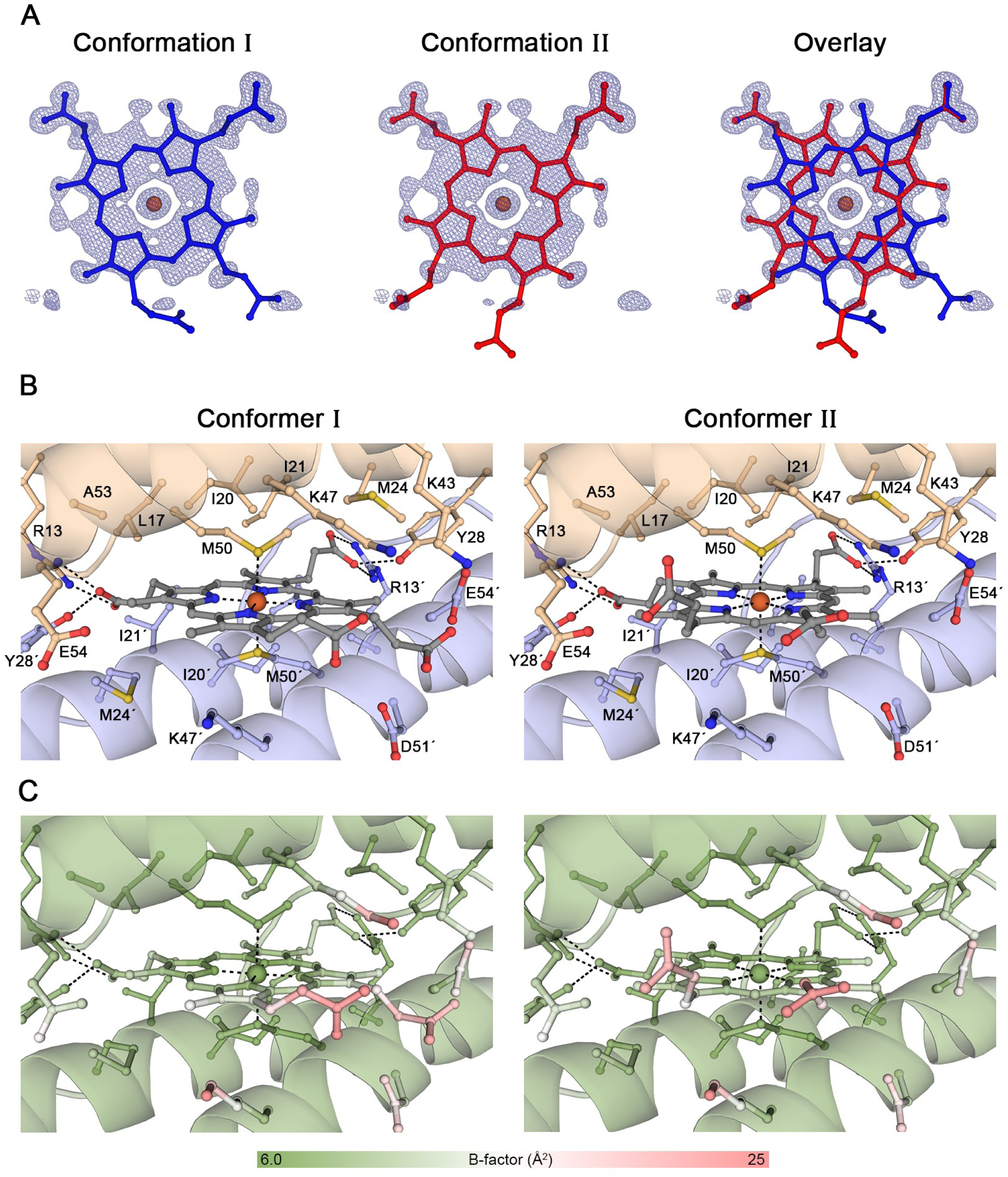
Coproheme coordination in *Mper*-mBfr. (A) Modelled coproheme with two alternative onformations and their overlap. The 2*F*_o_-*F*_c_ map contoured to 1.0 σ is represented as a green mesh. (B) Close-ups of the heme-binding pocket showing conformers independently. Beige and blue coils represent two different mBfr monomers. Oxygen, nitrogen, and sulfur are coloured in red, dark blue, and yellow, respectively. (C) Close-ups of the heme-binding pockets from the same view as (B) with a colour gradient reflecting the B-factors for each atom.

The coproheme is positioned within a hydrophobic pocket, where its two internally facing propionate arms are stabilised by hydrogen bonds with Arg13 and Tyr28 from both amino acid chains in the dimer (Fig. 4B), an interaction similar to that observed in *Dd*-Bfr. In contrast, the two solvent-exposed propionate arms extend toward the interior of the spherical assembly, but their lack of clear electron density reveals high flexibility, supported by the B-factor profile (Fig. 4C). The coproheme is accessible only from the interior and remains shielded from the external solvent (Fig. 3), a feature consistent with other bacterioferritins. Interestingly, in *Dd*-Bfr, the outward-facing propionate arms are not solvent-exposed but are further stabilized by interactions with C-terminal residues (13).

### The diiron ferroxidase centre is redox active *in crystallo*

The active site of *Mper*-mBfr conserved the canonical features of a bacterioferritin: two iron atoms coordinated by four glutamate residues (Glu16, Glu49, Glu91 and Glu124) and two histidine residues (His52 and His127) within the characteristic four-helix bundle (Fig. 5). Additionally, the Glu16 and Glu91 are stabilised by hydrogen bonding with Tyr23 and Tyr98. Each iron atom is hexacoordinated by the mentioned residues and a water molecule (Fig. 5), and separated by a distance of 3.9 Å, typical for a reduced ferrous state (13).

**Fig. 5.**
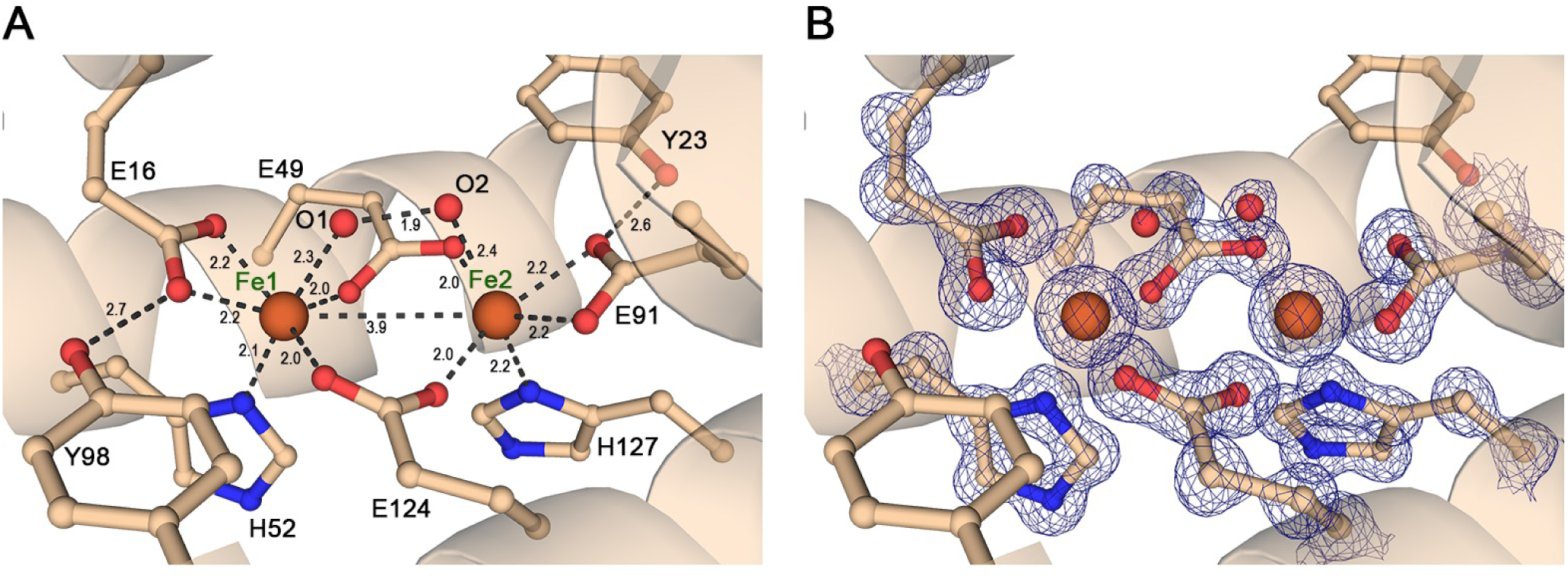
Diiron ferroxidase centre of *Mper*-mBfr. (A) Close-up of the ferroxidase centre showing amino acid residues (balls and sticks) involved in iron coordination within the four-helix bundle in light orange cartoon. Distances are displayed in Å. (B) Same view as (A) showing the amino acid residues involved in iron coordination, including the 2*F*_o_-*F*_c_ map in green contoured to 2.5 σ. Carbon, oxygen, nitrogen and iron are coloured as white, red, dark blue, and orange, respectively. Waters O1 and O2 are in two alternative conformations.

To investigate whether the ferroxidase centre of *Mper*-mBfr is redox active, protein crystals were subjected to an oxidation-reduction cycle. *In crystallo* spectrophotometry was performed to corroborate structural alterations induced by the oxidation from O_2_ exposure and back-reduction provoked by the dithionite agent. A control crystal representing the as-isolated state exhibited a spectrum with sharp peaks at 521 and 551 nm under cryogenic conditions, which was comparable to the solution spectrum at room temperature, confirming that the protein was in its reduced state, as previously observed (Figs. 5 and 6A). Upon oxidation by ambient air, a decrease in absorbance was observed. The decrease was more pronounced at 551 nm than at 521 nm. The latter is also getting wider. A second O_2_-exposed crystal that underwent the same colour change could be reduced back by soaking in dithionite, which restored the spectrum to its as-isolated state (Fig. 6A).

**Fig. 6.**
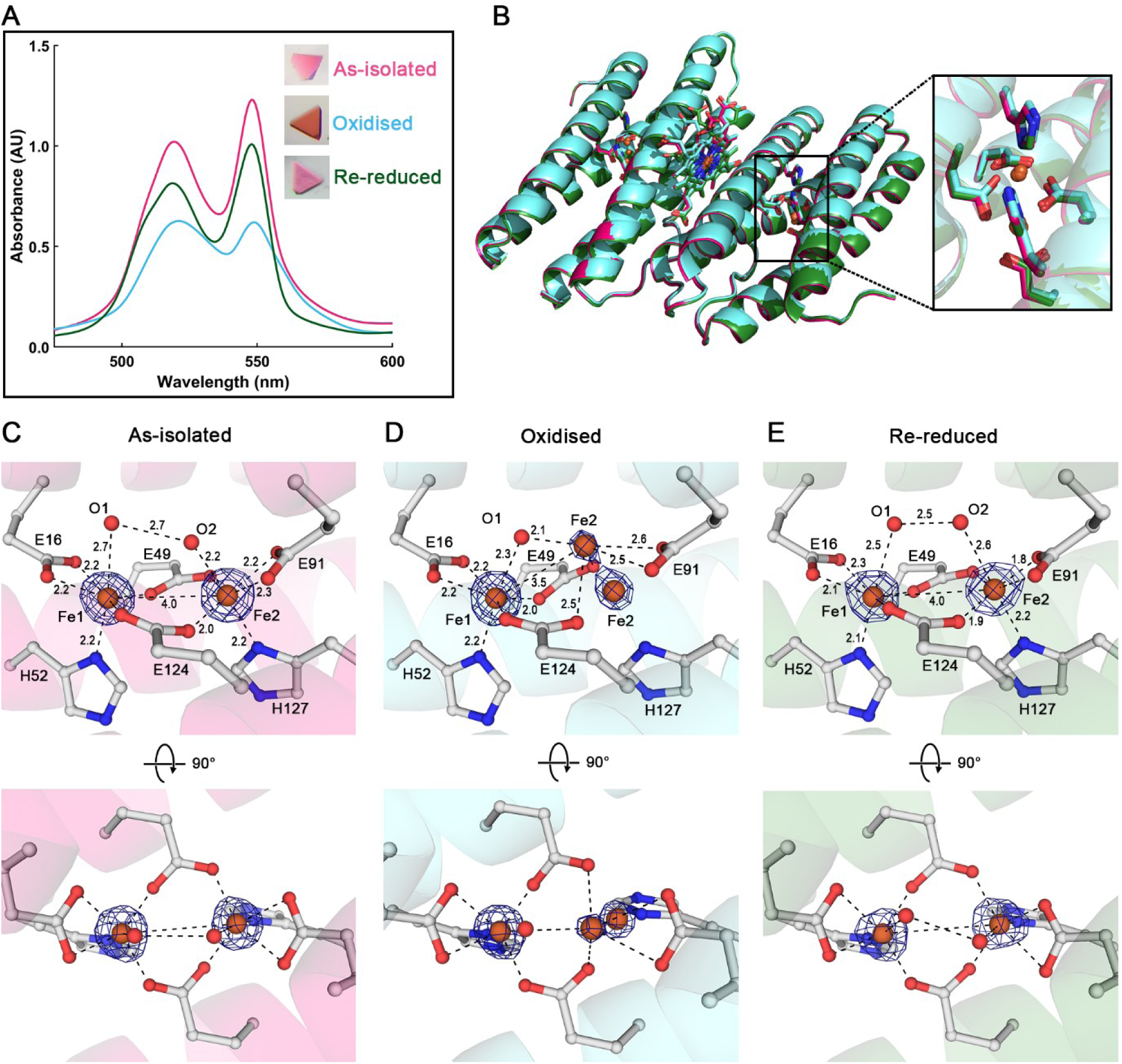
Redox cycling of *Mper*-mBfr crystals. (A) *In crystallo* spectrum of the as-isolated crystal, oxidised upon 10 minutes O_2_ exposure, and oxidised upon 10 minutes O_2_ exposure then reduced by soaking the crystals for 4 min 40 sec in 100 mM sodium dithionite. The inset displays photos of the corresponding crystals. (B) Overlay of a dimer of each structure, with as-isolated in pink, oxidised in cyan and re-reduced in dark green. The inset shows an overlay of the active site of all three conditions. (C-E) Zoom-in of the active site for all three conditions. Bottom panel displays a 90° rotation along the x-axis of the top panel. Distances between ligands are shown with dotted lines in Å. The dark blue mesh represents the 2*F*_o_-*F*_c_ map contoured to 7 σ. Ligands are represented as balls and sticks, with carbon, oxygen, nitrogen, and iron coloured in white, red, dark blue, and orange, respectively.

The three redox captured states did not show any major shift in the protein backbone with a root mean square deviation below 0.1 Å for around 132 Cα (Fig. 6B). A comparison of the ferroxidase centres (Fig. 6C-E) reveal a partial shift of Fe2 moving away from His127 and a slight twist of Glu49 in the oxidised state. The re-reduced state presents an identical coordination to the as-isolated state, with the Fe2 back into coordination with His127. A previous study on *Dd*-Bfr found Fe1 instead of Fe2 to be mobile upon redox cycling of protein crystals (13). This study also showed that redox cycling of *Dd*-Bfr in solution prior to crystallisation resulted in complete loss of Fe1 and conformational changes in the bridging glutamates. Such a dramatic effect might have been prevented by the crystal packing or might have happened if the crystals reached the fully oxidised state. A state that might have led to a loss of diffraction upon protein reorganisation.

### Phylogenetic analysis and classification of *Mper*-mBfr

To determine the phylogenetic placement of *Mper*-mBfr within the ferritin superfamily, we searched for homologous proteins (>30% identity and <10^−5^ e-value) and constructed a phylogenetic tree of the 81 closest related protein sequences alongside experimentally validated ferritin family members (Fig. 7A). The resulting tree revealed that *Mper*-mBfr does not cluster within the established clades of ferritins, bacterioferritins, Dps, or DpsL.

**Fig. 7.**
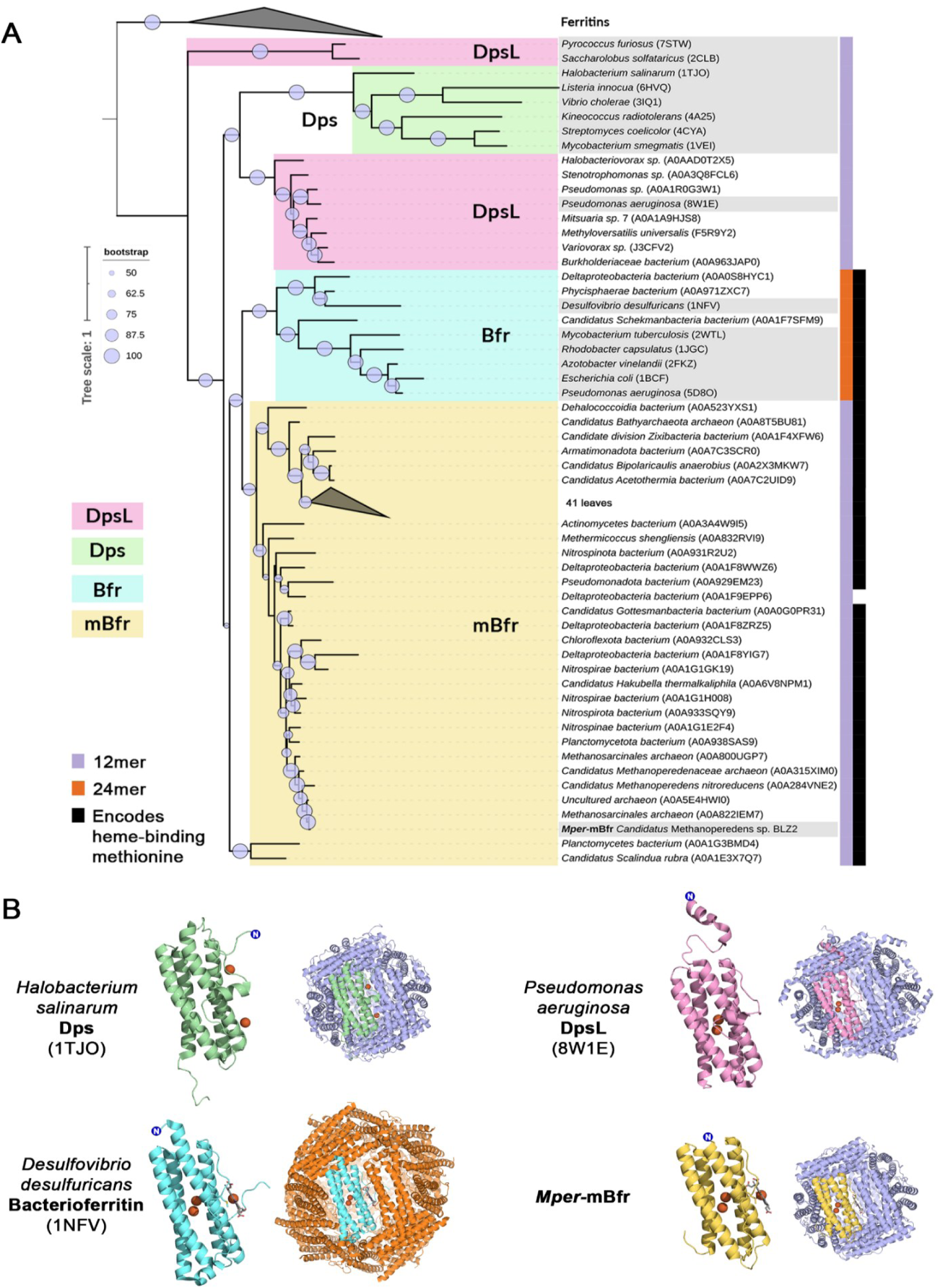
Phylogeny of mini-bacterioferritins. (A) Phylogenetic tree of amino acid sequences closely related to *Mper*-mBfr together with sequences from solved protein structures (highlighted in grey). The clades in pink, green, cyan, and yellow represent Dps-like proteins (DpsL), Dps, bacterioferritins (Bfr) and mini-bacterioferritins (mBfr), respectively. Multimerisation of the proteins was predicted with ColabFold and is indicated on the right as purple for 12-mers or orange for 24-mers. The presence of a heme-binding methionine is indicated in black. Bootstrap values in the range of 50-100 are indicated with purple circles. UniProt or RCSB PDB accession numbers are indicated in brackets. (B) Cartoon representation of four different types of ferritin-like proteins, including a monomer and its multimer. Colours of the monomers represent the different clades in the phylogenetic tree, and the colour of the additional subunits represents their multimer (purple for 12-mers and orange for 24-mers).

Since sequence identity alone did not provide insight into the quaternary structure of proteins in the phylogenetic tree, we explored whether protein structure prediction models could distinguish between 12-and 24-mer assemblies. Using ColabFold with an input of 12 copies of the amino acid sequence, we obtained structural predictions that formed either complete spheres or hemispheres (Supplementary Table S2, Supplementary Fig. S2). To further validate this approach, AlphaFold3 was used with 24 copies of the amino acid sequences, which resulted in the prediction of either two 12-mers or a single 24-mer assembly. The predicted structures were additionally analysed for methionines positioned at homodimeric interfaces that are suitable for heme binding. These structural predictions provided a basis for classifying phylogenetically related sequences according to their quaternary structure, offering a complementary approach to sequence-based classification.

This method identified that out of the 81 closest related protein sequences, 71 proteins, across a wide range of microorganisms, were predicted to fold into 12-mer spheres while also encoding a methionine for heme binding. These proteins were classified as mini-bacterioferritins (mBfr). Additionally, out of the 81 closest related proteins, seven were predicted to adopt a fold resembling a DpsL, while three aligned structurally with bacterioferritins.

A comparison of the monomer structures of Dps, DpsL, Bfr, and mBfr reveals that mBfr has the simplest composition, consisting solely of a four-helix bundle as its core structure. In contrast, other ferritin family proteins contain additional helices and flexible extensions (Fig. 7B). Dps and DpsL feature an additional helix positioned perpendicular to the four-helix bundle, located within the loop connecting helices B and C, at the position where (m)Bfrs bind heme. Some Dps and DpsL proteins, such as *Pa*-DpsL, possess additional N– or C-terminal tails associated with DNA-binding or endonuclease activity (21,34). In ferritins and bacterioferritins, an additional C-terminal E-helix extends from the four-helix bundle, forming the structural core of the four-fold symmetry axis in 24-subunit assemblies.

To identify sequence-level features that influence mBfr folding, we aligned the protein sequences of our predicted structures with the sequence of experimentally validated ferritin family proteins found in the RCSB PDB (Fig. 8A). The multiple sequence alignment of mBfr was then mapped onto a dimer structure using ConSurf (35). The ferroxidase centre, consisting of Glu18, Glu51, His54, Glu93, Glu126, and His129 (numbering shifted +2 relative to the *Mper*-mBfr sequence), appears to be conserved among DpsL, mBfr and Bfr.

**Fig. 8.**
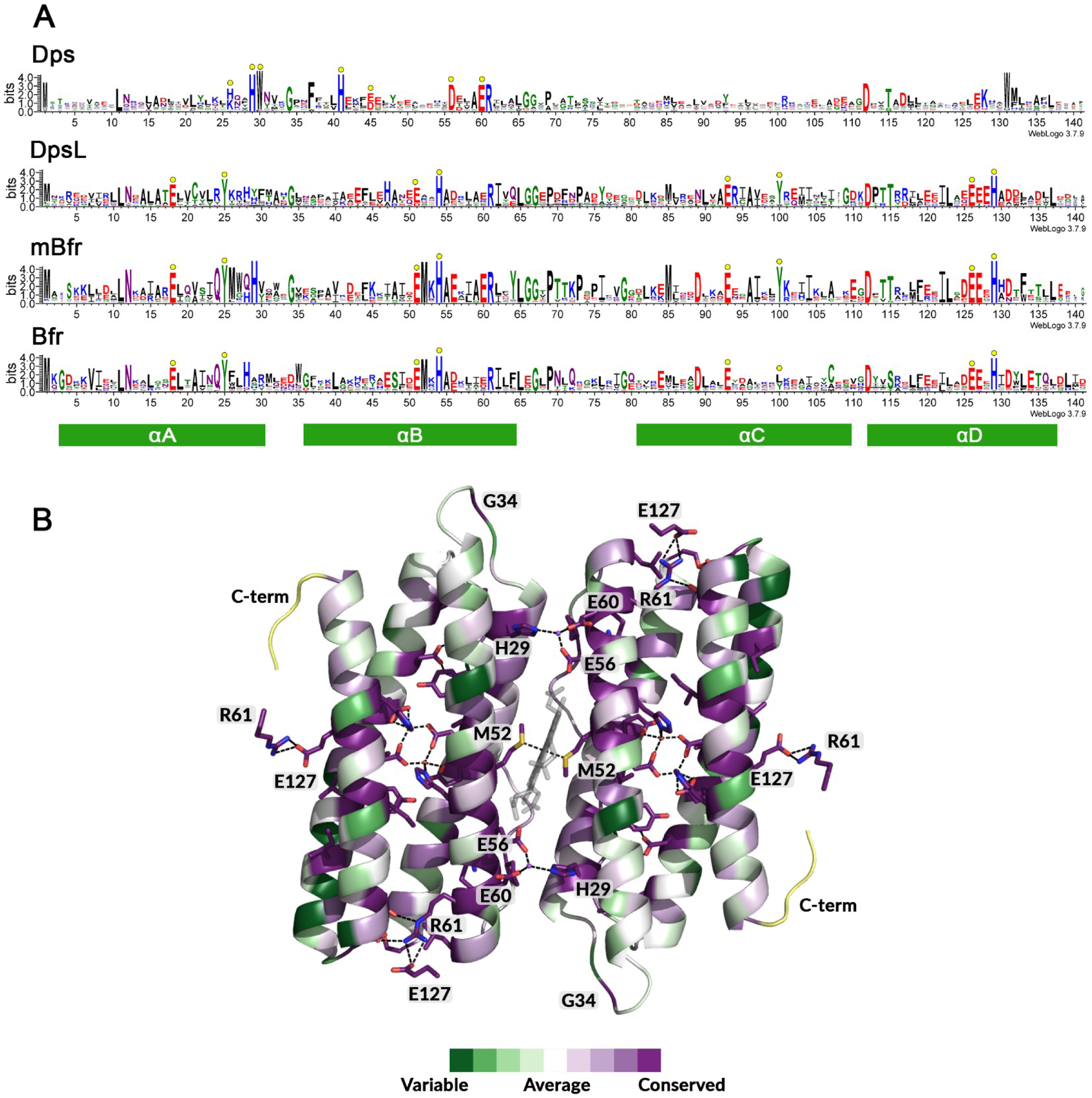
Multiple sequence alignments of ferritin-family proteins. (A) Multiple sequence alignments represented with WebLogo of Dps, Dps-like proteins (DpsL), mini-bacterioferritins (mBfr) and bacterioferritins (Bfr). Residues that are part of the active site are indicated by yellow circles. (B) ConSurf cartoon representation of the mBfr multiple sequence alignment showing conserved residues that interact with other amino acid chains. Regions with insufficient data are coloured yellow. Amino acid residues are numbered according to the multiple sequence alignment of mBfrs (+2 compared to *Mpe*r-mBfr). Ligands are represented as balls and sticks with oxygen, nitrogen and iron coloured in red, blue and orange, respectively.

We then examined conserved residues involved in inter-subunit interactions. The heme-binding methionine is strictly conserved in mBfr, mostly present in Bfr, and absent in other ferritin family members. Beyond methionines, a combination of three highly conserved residues, His29, Glu56, and Glu60, is present at the dimer interface. In the mBfr crystal structure, these residues coordinate a cation (tentatively modelled as a sodium) that links the monomers together, with His29 showing particularly strong conservation among mBfr proteins. Another key pair of residues, Arg61 and Glu127, form a salt bridge stabilizing dimer-dimer contacts. However, since these residues are also conserved in DpsL, Bfr, and partially in Dps, they do not provide a distinguishing feature for mBfr.

Another interesting residue is Gly34, located in the loop between helices A and B, which appears to be conserved exclusively in 12-mers Dps, DpsL, and mBfr. Taken together, the conservation of His29, Gly34, Met52, and an absence of N– and C-terminal extensions may serve as a useful signature for identifying new members of the mBfr group.

To contextualize these findings, we compared the defining features of major ferritin subfamilies: classical ferritins, bacterioferritins, Dps, DpsL, and the newly defined mBfr (Table 2). Among the family members, mBfrs stand out by its streamlined architecture, composed solely of a four-helix bundle, and their consistent 12-mer assembly with a strictly conserved heme-binding methionine. In contrast, other ferritin-like proteins show additional helices or tails associated with DNA binding. These distinctions emphasize the unique structural and functional identity of mBfrs within the broader ferritin superfamily and provide a framework for classifying newly discovered ferritin-like proteins.

**Table 2.** Overview of the different ferritin family members.

|  | **Ferritin** | **Bacterioferritin** | **Dps** | **DpsL** | **Mini-bacterioferritin** |
| --- | --- | --- | --- | --- | --- |
| **Ecological distribution** | Universal | prokaryotes | prokaryotes | prokaryotes | prokaryotes |
| **Multimerisation** | 24 | 24 | 12 | 12 | 12 |
| **Heme incorporation** | No | Yes, inter-subunit | No | No | Yes, inter-subunit |
| **DNA binding** | No | No | Yes, non-specific | Yes, non-specific (in most cases) | Unknown |
| **Diiron site type** | Canonical | Canonical | Inter subunit | Canonical | Canonical |
| **Primary function** | Iron storage & mobilisation | Iron storage & mobilisation | Stress response & DNA protection, limited iron storage | Stress response & DNA protection, iron storage | Unknown |

## Discussion

Universally conserved across living organisms, ferritins have been shown to be more than an iron storage, with physiological functions that could be expanded to microbial stress response and DNA protection. Here, we expand our understanding of ferritin natural diversity by isolating and characterising mini-bacterioferritins, the first member of a previously unrecognised group within the ferritin family.

The assembly of *Mper*-mBfr into a 12-mer quaternary structure, combined with the incorporation of coproheme in the same fashion as bacterioferritin, makes this ferritin-like protein unique. The only examples of 12-mer ferritins binding an heme are Dps from *Porphyromonas gingivalis* (*Pg*-Dps) and DpsA from *Synechococcus elongatus* (*Se*-Dps). In *Pg*-Dps, the heme is bound in a completely different way via a single surface-exposed cysteine and likely serves as a mechanism for heme storage and protection against heme toxicity (36). Regarding *Se*-Dps, the protein was found to bind heme *b* upon addition, though no specific binding mechanism was identified (37) and its sequence revealed no methionine residues positioned for heme coordination.

The incorporation of heme ligands facilitates the reduction of mineralised Fe^3+^ in the protein core to soluble Fe^2+^, as proposed in bacterioferritins (12). This heme is typically a heme *b* (33). However, in the case of *Mper*-Bfr, the ligand was identified as a coproheme, an intermediate of the coproporphyrin-dependent (CPD) heme *b* biosynthesis pathway found in archaea (38,39). Proteins that bind coproheme as a cofactor are rare and mainly limited to enzymes directly involved in the CPD pathway. Interestingly, the only other known example of a protein utilising coproheme as a cofactor is *Dd*-Bfr, which binds coproheme in a similar alternative dual conformation as *Mper*-mBfr (13,14). Redox titration of *Dd-*Bfr revealed an unusually high redox potential of +140 mV, contrasting with other bacterioferritins that typically exhibit redox potentials below −200 mV (28). This unique property raises intriguing questions about the function of coproheme in these proteins, particularly whether its presence influences electron transfer dynamics or interactions with physiological electron donors. The significantly higher redox potential suggests that coproheme may confer distinct reactivity compared to heme *b*, potentially affecting how these bacterioferritins participate in iron metabolism or oxidative stress responses. The reason for using coproheme instead of heme *b* remains unclear but may reflect specific adaptations to anaerobic environmental conditions. The heme identity in other mini-bacterioferritins is presumably different from coproheme, as the Arg13 stabilising the two propionate groups can be substituted by an Ile (Fig. 8A).

A phylogenetic analysis revealed that *Mper*-mBfr is part of a larger group of proteins characterized by a 12-mer assembly that can coordinate a heme at the dimer interface. Except for two outliers, all proteins predicted to be mini-bacterioferritins cluster into two distinct subclades and are found in both bacteria and archaea. Previous studies suggest that the 12-mer DpsL from *Saccharolobus solfataricus* and *Pyrococcus furiosus* may resemble an ancient common ancestor from which 24-mer bacterioferritins evolved (3,5). Although mini-bacterioferritins seem closer related to bacterioferritins than to DpsL, the structure of the last common ancestor shared by these groups remains unclear. Our sequence and structural analyses support the hypothesis that mini-bacterioferritins retain an ancestral 12-mer configuration, which may have predated the divergence of ferritin family proteins, and subsequently acquired heme-binding properties. Alternatively, DpsL may have lost its heme-binding ability during evolution.

The structural transition between 12– and 24-mer remains an open question. One study tested whether the C-terminal helix E, present in bacterioferritins and ferritins but absent in Dps, plays a defining role in oligomerisation. When helix E of *E. coli* Bfr was fused to *Ec*-Dps, the quaternary structure increased in size, but helix E flipped outward, and the protein still assembled as a 12-mer (40). In *Mycobacterium smegmatis*, a single Phe47Glu mutation was found to shift the oligomeric state to a 24-mer assembly, though only in crystallised form and not in solution (41). These findings suggest that while specific structural elements influence multimerisation, additional factors likely contribute to determining whether a ferritin-family protein assembles as a 12-mer or 24-mer.

The physiological function of *Mper*-mBfr remains speculative at this stage. Based on observed shifts in its active site upon oxidation, *Mper*-mBfr likely serves as an iron-storage protein, similar to *Dd-*Bfr (13). Additionally, it may contribute to oxygen protection, resembling the role of bacterioferritin in the anaerobe *Nitratidesulfovibrio vulgaris* (formerly *Desulfovibrio vulgaris*), which has been shown to mitigate oxidative stress (42). A study on salt stress in ‘*Ca.* Methanoperedens sp.’ Vercelli Strain 1 reported high expression of a 139-amino-acid ferritin-like protein with 87% sequence identity to *Mper*-mBfr, which protein modelling predicts to be a mini-bacterioferritin (43)(Supplementary Fig. S3). Interestingly, this protein was downregulated upon salt stress, suggesting a potential regulatory role in response to environmental changes.

The genome of ‘*Ca.* Methanoperedens sp.’ BLZ2 encodes four additional ferritin-like proteins, including two predicted rubrerythrins and two four-helix bundle proteins characteristic of the ferritin family (Supplementary Table S1, Supplementary Fig. S1). Based on the predicted tertiary structure of these two four-helix bundle ferritin-like proteins, it is difficult to assign them to any known ferritin subfamily, and modelling could not reliably predict their multimeric assembly. This genetic redundancy raises questions about the specific roles of these proteins and how they interact within the organism’s iron metabolism and stress response. To fully understand the physiological role of *Mper*-mBfr, *in vivo* studies are now needed to investigate its regulation, function, and potential interactions with other ferritin-like proteins in ‘*Ca.* Methanoperedens’. Genetically editable microbes harbouring mini-bacterioferritin could represent an attractive alternative.

More broadly, this study investigated ferritins beyond well-studied model organisms, showcasing the rich and largely untapped structural diversity of ferritin-like proteins in nature. *Mper*-mBfr could represent an example of evolutionary diversity driven by niche-specific adaptations, where the incorporation of coproheme (a rare cofactor) and a 12-mer quaternary structure distinguishes it from other well-studied ferritins and bacterioferritins. By tapping into organisms that are challenging to cultivate and from non-isolated lineages, we provided a new perspective on the ferritin family and their evolutionary path, highlighting the need to explore the natural microbial reservoir.

## Materials and Methods

### Bioreactor cultivation and lysis

Granular biomass highly enriched in ‘*Ca*. Methanoperedens’ BLZ2 (20-40%) was obtained from a sequencing fed batch bioreactor as described previously (44). About 30 g of anoxically stored biomass was defrosted in lukewarm water. Under a N_2_/CO_2_ (90:10) atmosphere, biomass was resuspended in a total volume of 90 mL IEC A buffer (50 mM Tris/HCl pH 8.0, 2 mM dithiothreitol (DTT)). The suspension was sonicated (30 sec, 75% power, 4 cycles, 1 min breaks, SONOPULSE Bandelin) to disrupt the granules, followed by five rounds of French Press at approximately 1,000 PSI (6.895 MPa). To prevent oxygen contamination, the French Press cell was flushed with N_2_ and washed twice with anoxic IEC A buffer. Cell debris was removed by centrifugation (45,000*g*, 30 min at 18 °C). Supernatant was sonicated again (30 sec, 75% power, 3 cycles, 1 min breaks) to reduce viscosity and subsequently subjected to ultracentrifugation (100,000*g*, 1.5 h at 4 °C, rotor 70Ti, Beckman Coulter) to prepare a soluble protein extract.

### Mini-bacterioferritin purification

Proteins were purified under anoxic conditions in an anaerobic tent with a N_2_/H_2_ (97:3) atmosphere at 20 °C under yellow light. The soluble extract (90 mL, 7.8 mg mL^-1^) was divided in two and loaded directly on 2 x 5 mL HiTrap Q HP columns (GE, Healthcare, Munich, Germany) pre-equilibrated with IEC A buffer. The loaded column was washed with IEC A buffer and proteins were eluded with a 0-65% gradient of IEC B buffer (50 mM Tris/HCl, pH 8.0, 1 M NaCl, and 2 mM DTT) in 80 min at 2 mL min^-1^ flow rate. 2 mL fractions were collected over the elution. Proteins were tracked using multi-wavelength absorbance monitoring (λ 280, 424 and 550 nm) in combination with denaturing PAGE. Fractions of interest were pooled and diluted 1:1 with HIC B buffer (25 mM Tris/HCl pH 8.0, 3 M (NH_4_)_2_SO_4_, and 2 mM DTT), filtered (0.2 µm nitrocellulose, Sartorius, Germany), and loaded on a 1.622 mL Source 15PHE 4.6/100 PE column (GE Healthcare, Munich, Germany) pre-equilibrated with HIC B buffer. Proteins were eluted using a 66 to 0% gradient of HIC B buffer mixed with HIC A buffer (25 mM Tris/HCl pH 8.0, and 2 mM DTT) in 70 min at 0.7 mL min^-1^ and collected in 0.5 mL fractions. Fractions of interest were pooled again and concentrated using a centrifugation concentrator (10 kDa cut-off, Vivaspin, Sigma-Aldrich, location). The final purification step included size exclusion chromatography on a Superdex 200 Increase 10/300 GL (GE Healthcare, Munich, Germany) in 25 mM Tris/HCl pH 8.0, 10% v/v glycerol and 2 mM DTT with a flow rate of 0.4 mL min^-1^, where *Mper*-mBfr showed a elution volume of 11.2 mL. This last step was skipped for the purification that yielded crystals for the redox cycle experiment. The final fractions of interest were concentrated and directly used for crystallisation or stored anoxically under N_2_ atmosphere at −80 °C.

### Protein identification by mass spectrometry

*Mper*-mBfr was identified using matrix-assisted laser desorption ionization time-of-flight mass spectrometry (MALDI-TOF-MS) as previously described (45). A spectrum range of 450-3000 *m/z* was recorded on Microflex LRF MALDI-TOF (Bruker, Billerica, MA, USA). Spectra were analysed using BioTools software (version 3.2, Bruker Life Sciences) linked to MASCOT search engine (Matrix Science Ltd., London, UK) loaded with the ‘*Ca.* Methanoperedens sp.’ BLZ2 proteome (NCBI:txid2035255, (23)). Peaks were selected with a minimal mass difference of 0.3 Da, a signal-to-noise threshold of 3 and a quality factor threshold of 20%. MASCOT search parameters included one trypsin miscleavage, global modifications of carbamidomethylated cysteines, variable modification of oxidised methionines, and a mass deviation of 0.2 Da. Eight observed peaks correlated with predicted *Mper*-mBfr peptides, resulting in a MOWSE score of 70.

### Heme identification by mass spectrometry

Heme was extracted from *Mper*-mBfr using a method from (46). To obtain enough hemes, several protein solutions (20-100 μL, 5-42 μg mL^-1^) were combined. 1 mL acetonitrile:HCl (ACN 0.8:0.2 HCl 1.7 M) was added to up to 50 μL protein solution and incubated on a shaker at room temperature for 20 min. Then, 250 μL saturated MgSO_4_ and 25 mg NaCl was added, and the solution was incubated on a shaker at room temperature for 5 min. This solution was then centrifuged (2500 *g* for 5 min) to obtain phase separation. The organic top layer containing the hemes was then extracted and stored at 4 °C.

The extract was analysed using a Agilent 1290 Infinity II LC system coupled to a 6546 Quadrupole Time of Flight mass spectrometer. A volume of 0.5 μL of the heme extract was injected onto a Poroshell 120 column (EC-C18, 2.1X50mm, 1.9μm; Agilent) maintained at 25 ^°^C, followed by elution with a gradient of mobile phase A (0.2% formic acid in water) and B (0.2% formic acid in acetonitrile) at a flow of 0.3 mL/min. The gradient was as follows: 0-15 min: 10-100%B, 15-25 min: 100%B, 25-26 min: 100-10%B, followed by 3 min re-equilibration at 10%B. One minute after injection, the eluate was directed to the Q-ToF MS operated in the positive ionization mode. MS1 scans were collected for m/z 50-1200, at a scan rate of 4 spectra/s. MS2 fragmentation spectra were collected using the same setup, but collecting targeted MS2 scans of m/z 706.2 at collision energies of 20, 40, and 60. MS1 and MS2 spectra of the peak with m/z 706.2 were imported into SIRIUS v6.10 (47,48) for the prediction of elemental composition and structure. The elemental composition was determined using de novo prediction, limiting the number of nitrogens to 4 based on the porphyrin structure observed in the crystal structure.

### High-resolution clear native PAGE (hrCN PAGE)

The PAGE was prepared and run as in (49). The whole process was performed anoxically. Glycerol (20 % v/v final) was added to each sample, and 0.001 % (w/v) Ponceau S serves as a marker for protein migration. The electrophoresis cathode buffer contained 50 mM Tricine; 15 mM Bis-Tris, pH 7; 0.05 % (w/v) sodium deoxycholate; 0.01 % (w/v) dodecyl maltoside and 2 mM DTT. The anode buffer contained 50 mM Bis-Tris buffer pH 7, and 2 mM DTT. hrCN PAGE was carried out using a 5 to 15 % linear polyacrylamide gradient, and gels were run with a constant 40 mA current (PowerPac™ Basic Power Supply, Bio-Rad).

### Crystallisation

An initial screening using the sitting drop method was performed on a 96-well MRC 2-well polystyrene crystallisation plate (SWISSCI, Switzerland) at 20 °C under a pure N_2_ atmosphere. The plate was filled with 90 µL crystallisation solution (Wizard screen from Jena Bioscience, Germany) in the large reservoir and the purified protein solution concentrated to 4.4 mg/mL was spotted in the small reservoirs by mixing 0.5 µL purified protein with 0.5 µL crystallisation solution using an OryxNano robot (Douglas Instruments Ltd, UK). Once prepared, the plate was transferred to an anaerobic chamber containing a N_2_/H_2_ (97/3%) atmosphere for long-term storage. The crystal diffracting to atomic resolution was harvested from the original screening plate containing the following crystallisation solution: 20% v/v Jeffamine M-600 pH 7.0, and 100 mM HEPES pH 7.5. Prior to liquid nitrogen freezing, the crystal was soaked in the crystallisation solution supplemented with 25% v/v glycerol for a few seconds.

For the redox cycling experiment, crystals were obtained on a Junior Clover plate (Jena Bioscience, Germany) using 100 µL crystallisation solution in the reservoir composed of 20% w/v Polyethylene glycol 3,350 and 200 mM tri-Potassium citrate. 1 µl of purified protein solution at 22 mg/mL was mixed with 1 µl of the crystallisation solution. The crystal corresponding to the as-isolated state was directly harvested inside the anaerobic chamber, and the one corresponding to the oxidised state was exposed to ambient air for 10 minutes. For the re-reduced state, the crystal was exposed to ambient air for 10 minutes, then soaked in the crystallisation solution supplemented with 100 mM of freshly prepared sodium dithionite. All harvested crystals were soaked for a few seconds in the crystallisation solution supplemented with 20% v/v glycerol, except for the re-reduced state, where the crystal was soaked in the crystallisation solution supplemented with 20% v/v glycerol and 90 mM sodium dithionite.

### X-ray data collection and model refinement/validation

Data were collected (see Table 1) at the Swiss Light Source (SLS, beamline PXI-X06SA) and at the European Synchrotron Radiation Facility (ESRF, beamline BM07-FIP2) under cryogenic conditions (100 K). All data were integrated with *autoPROC* (50). All datasets were treated as isotropic, except for the atomic resolution dataset, which exhibited a slight anisotropy. The atomic resolution structure was solved by molecular replacement using *Phaser* from the *PHENIX* package (51) with an Alphafold2 model (52) of the *Mper*-mBfr monomer. The three other structures from the redox cycle were solved in the same way, with the atomic resolution model as a template.

All models were manually optimised with *Coot* (v0.9.8) (53). Refinement was performed with *Phenix* refine (v1.21.1-5286) (51) without applying non-crystallography symmetry. All models were refined by using a translation-libration screw except for the atomic resolution model in which all atoms were considered anisotropic. All models were refined by adding hydrogens in riding positions. Models only contain hydrogens on the protein chains and not on ligands and solvent. The final structures deposited to the PDB did not contain hydrogens except for the atomic resolution dataset. The different structures were validated by the *MolProbity* tool integrated with *PHENIX* (54).

### Structural analyses

The inner volume of *Mper*-mBfr, *Hs*-Dps, *Pa*-DpsL and *Dd*-Bfr was measured using the program HOLLOW (55), by modelling a cylinder in the inner cavity using a grid-spacing of 0.5 Å in the case of *Mper*-mBfr, *Hs*-Dps and *Pa*-DpsL or 1.0 Å in the case of *Dd*-Bfr and manual curation in PyMOL (v2.2.0, Schrödinger, LLC). PyMOL was further used for structural analysis and visualisation of protein structures.

### UV/visible spectrophotometry

UV-Vis spectra were recorded at a protein concentration of 2 mg mL^-1^ on a Cary 60 (Agilent Technologies, USA) in a stoppered anoxic quartz cuvette with a light pathlength of 1 cm. The oxidised spectrum was recorded after opening the stoppered cuvette and equilibrating with air for 10 min. The same sample was reduced again inside an anaerobic chamber with a N_2_/H_2_ (97/3%) atmosphere by adding freshly prepared sodium dithionite (100 mM stock solution) to a final concentration of 1 mM, and stoppering the cuvette. The spectrum was recorded after 5 min.

UV-Vis absorption spectra of *Mper*-mBfr crystals were collected at the *in crystallo* Optical Spectroscopy (*ic*OS) laboratory at the ESRF (56). Spectra were measured using a DH-2000-BAL deuterium-halogen lamp (Ocean Optics, Dunedin, FL) as the reference light source and a QE65 Pro spectrophotometer (Ocean Optics, Dunedin, FL).

Crystals were mounted in a standard crystallisation loop between the two reflective objectives of the *ic*OS setup and maintained at 100 K using an evaporating nitrogen cryostream (Oxford Cryosystems). The lamp was connected to the upper objective via a 200 µm-diameter fibre. The spectrometer was connected to the lower objective via a 400 µm-diameter fibre. Spectra were recorded with a 6 ms acquisition time and averaged over 10 scans. Spectra were prepared using the *ic*OS toolbox (57).

### Phylogenetic analysis

The ferritins amino acid sequences used to build the phylogenetic analyses in Fig. 7 were recovered from a database constructed by combining all sequences associated with InterPro IPR002024 (Bacterioferritin) and Pfam PF00210 (Ferritin-like domain) protein families. Duplicated sequences were removed using seqkit v.2.8.0 (function rmdup) (58) resulting in a total of 68990 unique protein sequences in the database. A DIAMOND v.2.1.6 (59) blastp search was done using the sequence of *Mper*-mBfr (UniProt A0A0P8A5A1) as reference. The top 1000 matches that respected a cut off of >30% identity and <E−05 e-value were used in the final analysis. Additionally, 15 ferritin family protein sequences with experimentally confirmed structures were obtained from the RCSB PDB. UniProt or RCSB PDB accession numbers of the proteins used in the alignment are displayed in brackets in Fig. 7. The sequences were aligned using MAFFT v7.397 (60) and regions in the alignment containing >50% gaps were removed using the web version of ClipKIT (61). The tree was constructed using IQ-TREE v2.3.6 (62), where the best model (LG+F+I+G4) was chosen according to the Bayesian Information Criterion (BIC) and 1000 bootstraps were executed. The final visualisation and annotation of the phylogeny were done in iToL v7.1 (63).

To predict the multimerisation of the proteins in the tree ColabFold (64) was used, giving 12 amino acid chains input per protein, using 1-5 models and 3 recycles per protein. The predicted models were analysed in PyMOL (v2.2.0, Schrödinger, LLC), showing multimers folded in either a perfect 12-mer sphere or a 12-mer hemisphere, indicating a likely 24-mer multimerisation. Examples of the predicted structures are shown in Supplementary Fig. S2, and the confidence matrices for all predicted structures are provided in Supplementary Tables S1 and S2.

To visualise the conserved residues, the Consurf server (35) was used with the atomic resolution *Mper*-mBfr structure together with the multiple-sequence alignment of all predicted mBfrs (Supplementary table S3) and otherwise default parameters.

## Acknowledgements

We would like to thank the Max Planck Institute for Marine Microbiology and the Max Planck Society for their continuous support. We thank Christina Probian and Ramona Appel of the Microbial Metabolism laboratory for their assistance and support, and Marie-Caroline for performing the initial crystallisation screen. We thank Huub op den Camp for his help with protein mass spectrometry. We acknowledge the SLS and ESRF synchrotrons for allocating beam time and the beamline staff of PXI-X06SA and BM07-FIP-2 for their advice and assistance during data collection. T.W. was supported by a European Research Council consolidator grant [Grant number 101125699] and the Max Planck Society. The initial crystallisation screening performed by an OryxNano robot was financed by the DFG priority programme 1927 Iron-Sulphur for Life [Grant number WA 4053/1-1]. This study was supported by the SIAM Gravitation grant funded by NWO [Grant number 024.002.002] and an NWO-VIDI Talent grant [Grant number VI.Vidi.223.012]. It was furthermore supported by the ERC Synergy Grant MARIX [Grant number 854088]. We acknowledge the French Biology/Health Panel Review Committee for the provision of synchrotron radiation beamtime at the ESRF (Grenoble, France) on beamline BM07-FIP2, supported by the French ANR PIA3 (France 2030) EquipEx+ project MAGNIFIX under grant agreement ANR-21-ESRE-0011. This work used the *ic*OS platform of the Grenoble Instruct-ERIC centre (ISBG; UAR 3518 CNRS-CEA-UGA-EMBL) within the Grenoble Partnership for Structural Biology (PSB), supported by FRISBI (ANR-10-INBS-0005-02) and GRAL, financed within the University Grenoble Alpes graduate school (Ecoles Universitaires de Recherche) CBH-EUR-GS (ANR-17-EURE-0003).

## Author contributions

Microbial cultivation was done by M.W. and C.W. Purification and crystallisation were performed by M.W. and T.W. X-ray data collection was done by S.E. and T.W. X-ray data processing, model refinement, and analysis were done by M.W. and T.W. *In crystallo* spectrophotometry and analysis were done by S.E. and A.R. Mass spectrometry was done by M.W. Phylogeny analyses were performed by M.W. and P.L. The manuscript was written by M.W. Manuscript edition and correction were done by all authors.

