## Supplementary Data for "Mini-bacterioferritins: Structural insight into a new type of ferritin-like protein from an anaerobic methane-oxidising archaeon"

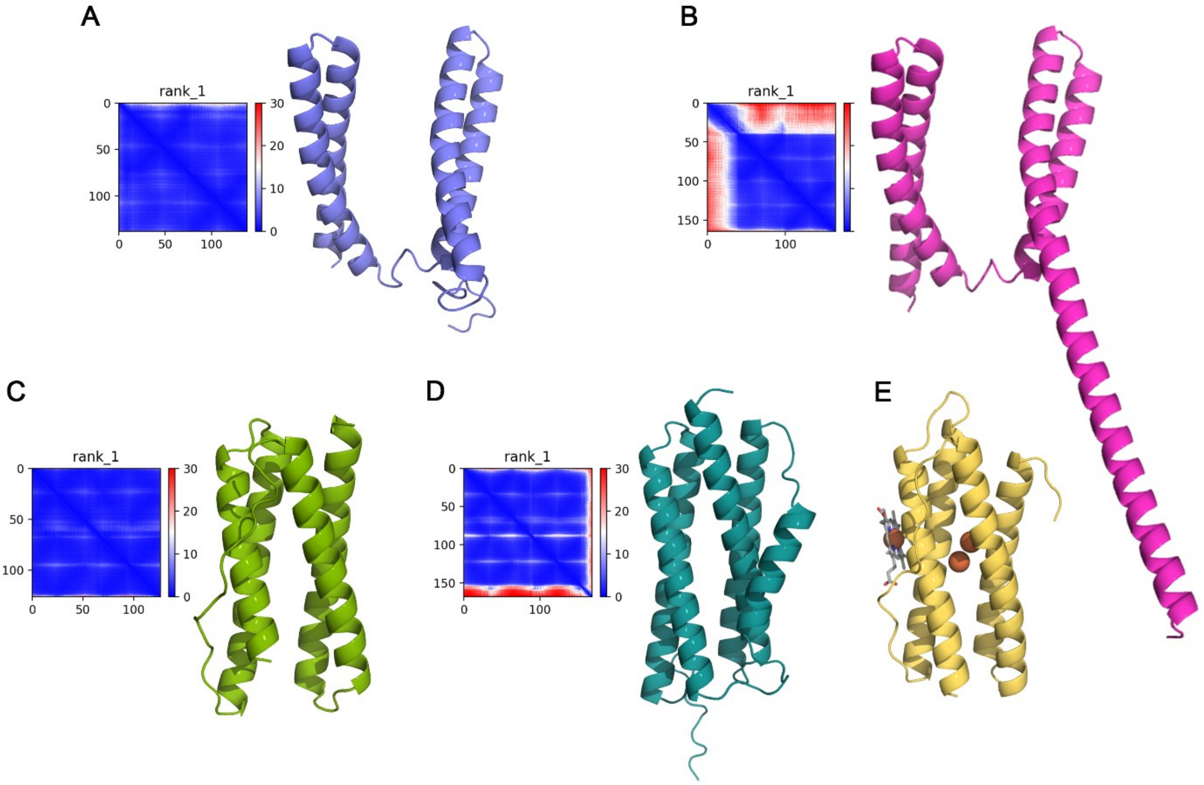


**Supplementary Fig. S1. AlphaFold models of ferritin-like proteins identified in '*Ca.* Methanoperedens sp.' BLZ2.** Each panel displays the Predicted Aligned Error (PAE) plot alongside the corresponding protein model in cartoon representation. (A) WP_097297487.1 (UniProt accession A0A6A2G1C8). (B) WP_176505107.1 (UniProt accession UPI0015968CAC). (C) WP_217993135.1 (UniProt accession A0A0P7ZGQ0). (D) WP_097298589.1 (UniProt accession A0A822J9T9). (E) Cartoon representation of *Mper*-mBfr.

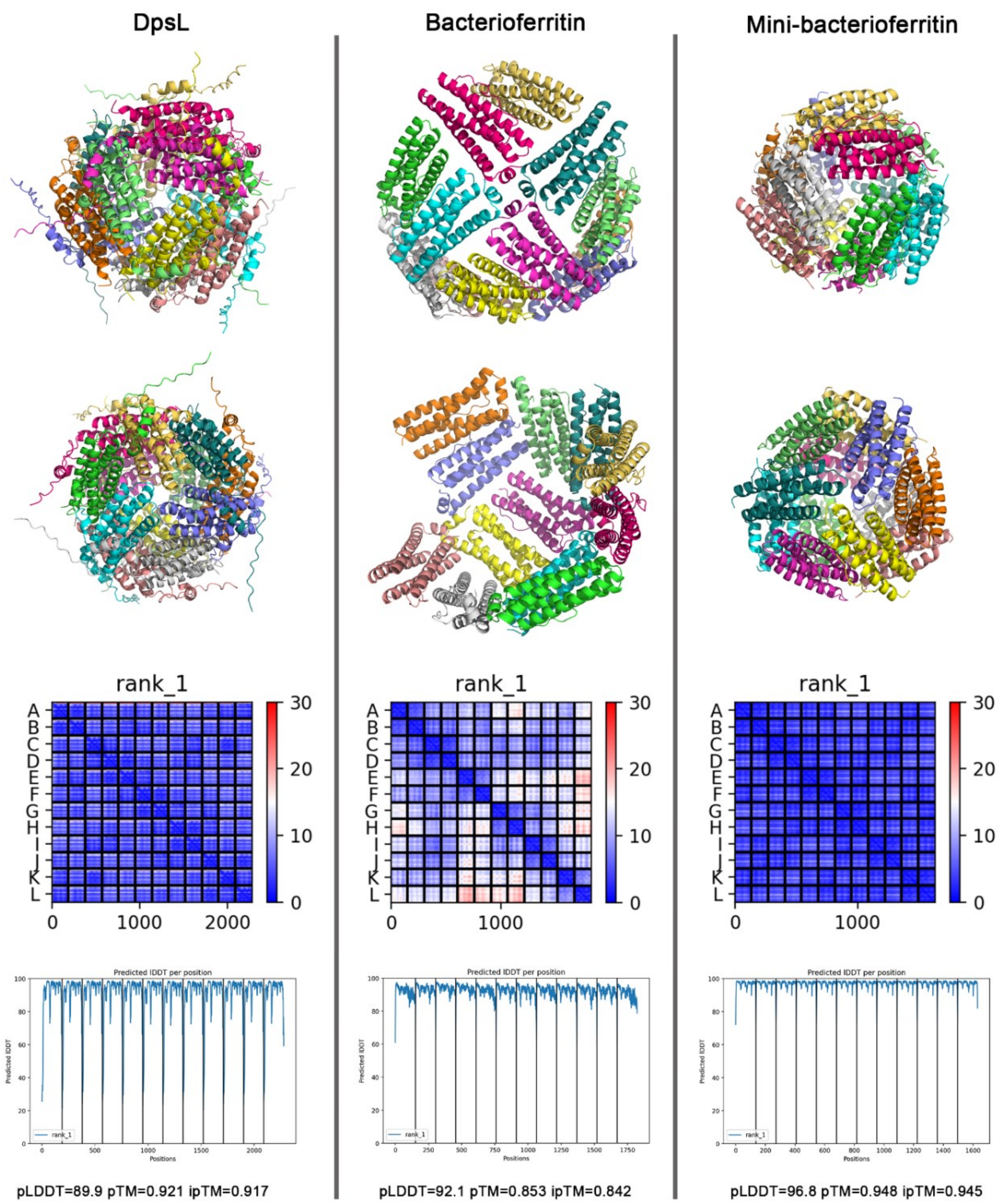


**Supplementary Fig. S2. Protein model representatives.** Protein models were generated using ColabFold with 12 copies as input to classify proteins in the phylogenetic tree (Fig. 7). This figure presents the three predicted types of ferritins: DpsL (left) from *Stenotrophomonas sp*. (UniProt accession code A0A3Q8FCL6), bacterioferritin (middle) from *Deltaproteobacteria* bacterium (UniProt accession code A0A0S8HYC1), and mini-bacterioferritin (right) from '*Candidatus* Scalindua rubra' (UniProt accession code A0A1E3X7Q7). From top to bottom, the models are shown in two different orientations using a cartoon representation, followed by the Predicted Aligned Error (PAE) plot and the per-residue pLDDT confidence score plot.

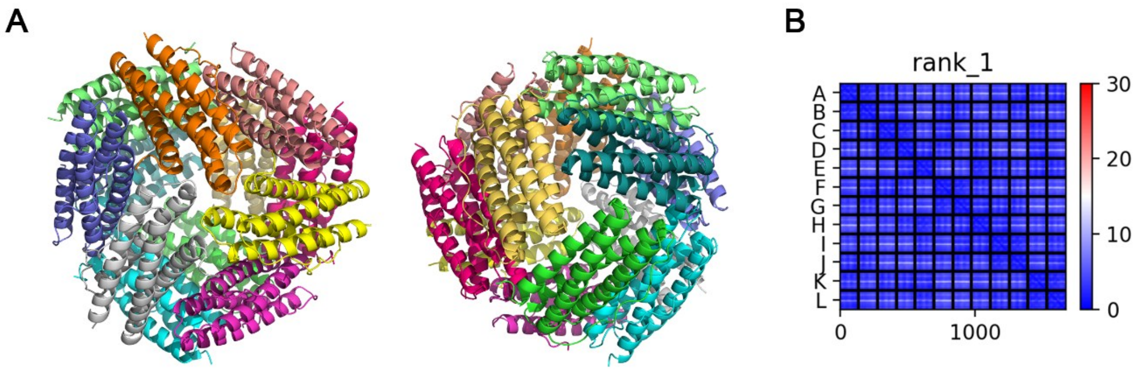


**Supplementary Fig. S3. Protein model of '*Ca.* Methanoperedens sp.' Vercelli mini-bacterioferritin.** (A) View of the protein from the C-terminal threefold axis (left) and N-terminal threefold axis (right). (B) Predicted Aligned Error (PAE) plot of the model.
